# Molecular basis of RNase I-mediated rRNA degradation regulated by ribosomes

**DOI:** 10.1101/2024.07.29.605612

**Authors:** Atsushi Minami, Takehito Tanzawa, Tomoko Miyata, Fumiaki Makino, Zhuohao Yang, Takashi Funatsu, Tomohisa Kuzuyama, Hideji Yoshida, Takayuki Kato, Tetsuhiro Ogawa

## Abstract

Protein biosynthesis is an energy-hungry intracellular process that requires the stringent regulation of ribosome abundance under environmental conditions. In response to stress, some active ribosomes are degraded while others, in bacteria, enter a hibernation state to protect against degradation. RNase I, a conserved T2 family ribonuclease in *Escherichia coli*, degrades ribosomal RNA to suppress biofilm formation, whereas it interacts with ribosomes. However, how and why RNase I binds to ribosomes remains elusive. Here, we show that hibernating ribosomes bind to RNase I and inhibit its activity, thereby promoting biofilm formation. We determined the cryo-electron microscopy structure of the hibernating ribosome complexed with RNase I. RNase I interacts with helix 41 of the 16S rRNA and the ribosomal protein uS14 in the head domain of the 30S subunit of the hibernating ribosome and positions its active centre away from helix 41, resulting in its catalytic inactivation. Hibernating ribosomes are protected from RNase I-mediated cleavage, and our *in vivo* and *in vitro* analyses revealed that RNase I targets dissociated large and small ribosomal subunits for rRNA degradation. These findings reveal a previously uncharacterized regulatory strategy that ribosomes modulate RNase I activity, ensuring both the preservation and timely degradation of ribosomes during environmental stress adaptation.

## Main

Ribosomes are sophisticated molecular machines that translate genetic information into proteins in all domains of life. Since protein biosynthesis consumes substantial energy, cells tightly regulate ribosome abundance in response to environmental conditions. In bacteria, under stress conditions such as nutrient starvation, heat shock, oxidative stress, or entry into the stationary phase, ribosome biogenesis and translation initiation are promptly suppressed^1^, and excess active ribosomes are degraded^2^.

Conversely, a subset of ribosomes is protected from degradation to ensure the rapid resumption of translation once favourable conditions return. One strategy for ribosome protection under stress is ‘ribosome hibernation,’ which preserves ribosomes in a dormant state. This mechanism also prevents unnecessary translation for suppressing energy consumption^3–6^. In many bacteria, ribosomes typically dimerize to form 100S ribosomes with the involvement of ribosome-binding hibernation protein factors. Several hibernation factors have been identified, including ribosome modulation factor (RMF), hibernation-promoting factor (HPF), ribosome-associated inhibitor A (RaiA), and the recently discovered Balon^3,7^. Ribosome hibernation is also reported in eukaryotes^8–11^.

Ribonucleases (RNases) regulate cellular RNA metabolism by degrading various types of RNA, including messenger RNA, transfer RNA, and ribosomal RNA. The degradation of rRNA is essential for eliminating excess ribosomes. Among many ribonucleases, RNase T2 is a crucial endoribonuclease that degrades rRNA in various organisms^12–14^ such as yeast RNase T2 (Rny1), which degrades rRNA during autophagy^15,16^. RNase T2 was first discovered in *Aspergillus oryzae*^17^ and later identified in a wide range of organisms. It cleaves single-stranded RNA non-specifically using two catalytic histidine residues^18^. The RNA cleavage mechanism consists of two steps: the phosphodiester bond cleavage by phosphate transfer, which forms an intermediate 2′,3′-cyclic nucleotide monophosphate (2′,3′-cNMPs), followed by hydrolysis to form the 3′-phosphate group. However, in some cellular contexts, cyclic intermediates accumulate because the second hydrolysis step is kinetically limited^19,20^.

In contrast to RNase T2 in eukaryotes^12–14,21–23^, little is known about its biological functions in bacteria. *Escherichia coli* RNase T2, also called RNase I, is a major ribonuclease that degrades rRNA, as in other species^24,25^. In addition, RNase I binds to the ribosomal 30S subunit^26–28^. Although RNase I is predominantly localized in the periplasm^26^, these findings suggest that a fraction of RNase I resides and functions in the cytoplasm. The molecular basis for this interaction and translocation remains unclear; however, exposure to mercuric ions triggers extensive RNA degradation by RNase I in *E. coli*, implying a role in stress-induced RNA decay^24^. Recently, it was shown that 2′,3′-cNMPs, generated by RNase I, serve as second messengers and inhibit biofilm formation^29^. Biofilms are aggregated communities of various microorganisms formed under nutrient limitations and stress^30^. Thus, biofilm formation requires a reduction in 2′,3′-cNMP levels, i.e., to inhibit RNase I activity under such conditions. However, how RNase I activity is regulated in response to environmental conditions remains unknown. To investigate the underlying mechanisms of this regulation, we focused on the interaction between RNase I and hibernating ribosomes, which accumulate under biofilm-forming stress conditions. In this study, we demonstrate that RNase I activity is inhibited by hibernating ribosomes both *in vitro* and *in vivo*, thereby promoting biofilm formation. To understand the structural basis of this inhibition, we determined the cryo-EM structure of the hibernating ribosome in complex with RNase I at a resolution of 2.7 Å. This revealed that RNase I specifically binds to the head domain of the 30S subunit. Moreover, our biochemical analyses explain the preferential association of RNase I with non-translating ribosomes and reveal that ribosome dissociation into subunits exposes rRNAs to cleavage by RNase I. Together, these findings elucidate the mechanism by which ribosomes dynamically regulate their vulnerability to ribonuclease attacks, offering new insights into the coevolution of ribosomes and RNA-degrading enzymes.

### RNase I-mediated rRNA degradation is suppressed in the *E. coli* strain accumulating hibernating ribosomes

In *E. coli*, a hibernating ribosome (100S ribosome) is formed by the dimerization of two 70S ribosomes, and the formation is promoted by the binding of HPF and RMF. The RaiA-deficient (Δ*raiA*) strain, lacking another hibernation factor RaiA, accumulates 100S ribosomes in the cytoplasm^31,32^ (Supplementary Fig. 1a). We first compared rRNA degradation among wild-type, RNase I-deficient (Δ*rna*), and Δ*raiA* strains under amino acid starvation. After 2, 4, or 24 h of starvation, total RNA was extracted and analysed by electrophoresis (Extended Data Fig. 1). At 2 h, rRNA degradation was largely suppressed in both wild-type and Δ*raiA* strains. By 4 h, substantial rRNA degradation occured in the wild-type strain, whereas rRNA was relatively preserved in the Δ*raiA* strain. This was supported by quantitative analysis using a Bioanalyzer, which confirmed that after 4 h of starvation, the wild-type strain retained 40% of 16S and 21% of 23S rRNA, whereas the Δ*raiA* strain preserved 76% and 49%, respectively (Fig. 1a). These results indicate that rRNA degradation is inhibited in the Δ*raiA* strain compared with that in the wild-type strain. However, when cultivation under starvation was continued for up to 24 h, extensive rRNA degradation occured in both strains, indicating that the protective effect in the Δ*raiA* strain is not sustained under prolonged starvation (Extended Data Fig. 1). Almost no degradation was observed in the Δ*rna* strain even after 24 h of starvation, confirming that RNase I is the primary ribonuclease responsible for rRNA degradation during starvation (Fig. 1a and Extended Data Fig. 1).

**Fig. 1.**
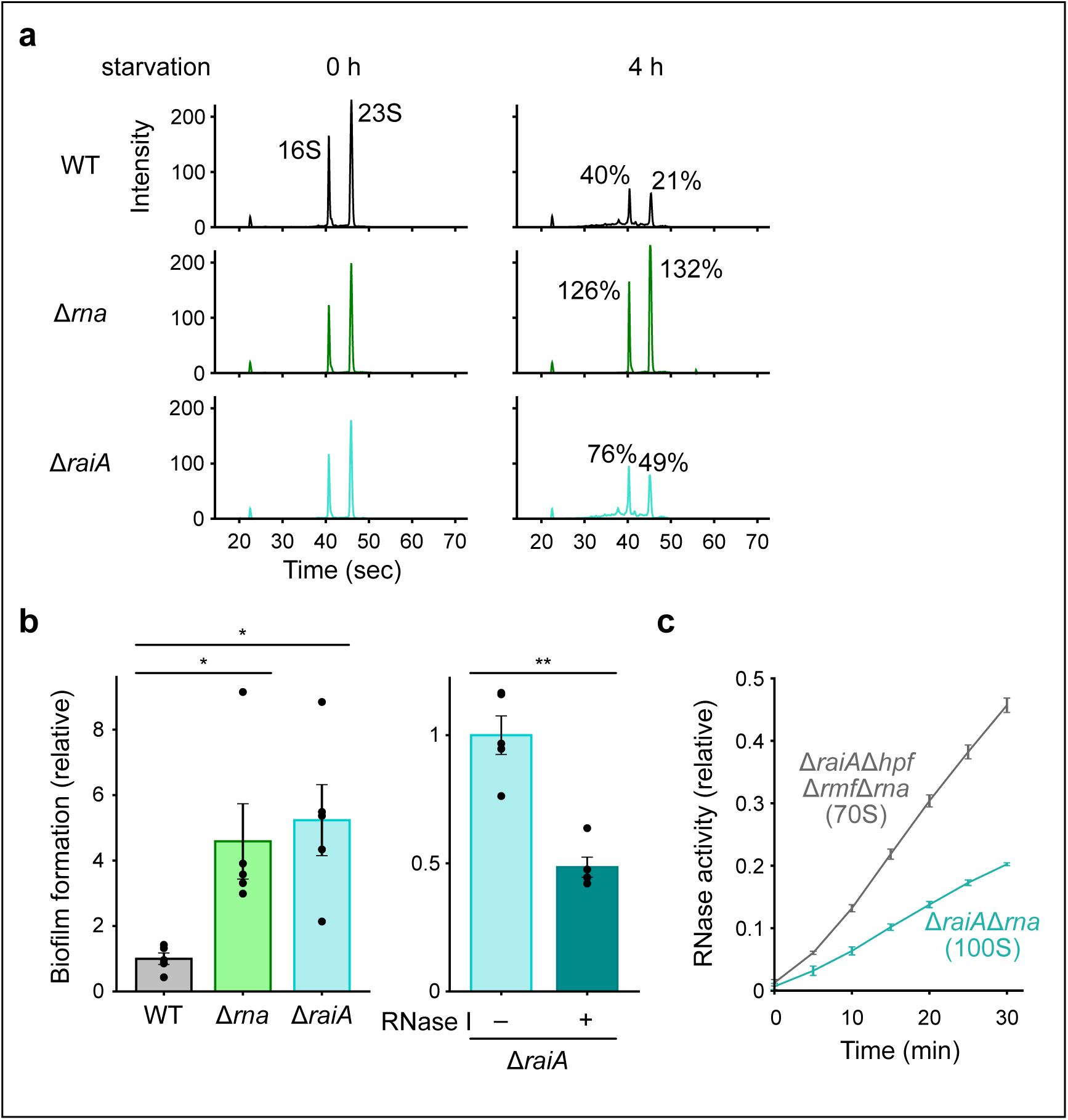
Suppression of RNase I-mediated rRNA degradation during starvation enhances biofilm formation in the Δ*raiA* strain. **a**, Quantification of total RNAs prepared from wild-type (WT), Δ*rna*, and Δ*raiA* strains during amino acid starvation. Samples are arranged from top to bottom as WT, Δ*rna*, and Δ*raiA*, and from left to right as 0 h and 4 h after starvation. **b**, (left) Biofilm-forming ability of indicated strains during amino acid starvation. (right) Biofilm formation in the Δ*raiA* strain with or without RNase I overexpression. **c**, Comparison of the inhibitory activity of 70S ribosomes (from Δ*raiA*Δ*hpf*Δ*rmf*Δ*rna* cells) and 100S ribosomes (from Δ*raiA*Δ*rna* cells) against RNase I. Data in **b** and **c** represent mean ± standard error from five and three independent experiments, respectively. Asterisks indicate statistically significant differences (*P < 0.05, **P < 0.01; Welch’s t-test).

### Inhibition of RNase I by hibernating ribosomes promotes biofilm formation

Based on the above results, we next measured biofilm-forming ability. In the Δ*raiA* and Δ*rna* strains, biofilm formation increased over 4-fold compared with that in the wild-type strain (Fig. 1b, left panel). In the Δ*raiA* strain, biofilm levels decreased upon RNase I overexpression (Fig. 1b, right panel), supporting our hypothesis that hibernating ribosomes inhibit RNase I, leading to increased biofilm production in the Δ*raiA* strain. To investigate how ribosome-associated RNase I activity is regulated, we examined its interaction with hibernating ribosomes *in vivo*. Since most RNase I is localized in the periplasm^26^, we extracted spheroplasts to remove periplasmic RNase I and isolated ribosomes from the resulting cytoplasmic lysate by ultrafiltration. Western blotting showed that the ribosome fraction from the Δ*raiA* strain contained more RNase I than that from the wild-type strain (Extended Data Fig. 2a). Further, higher RNase I activity was detected in the ribosome fraction from the Δ*raiA* strain than the wild-type strain (Extended Data Fig. 2b). These results suggest that hibernating ribosomes bind more readily to RNase I than 70S ribosomes. Then, hibernating ribosomes were tested for their ability to inhibit RNase I activity *in vitro*. Ribosomes were isolated from a Δ*rna* strain accumulating 100S ribosomes (Δ*raiA*Δ*rna*) and from a strain lacking all hibernating factors (Δ*raiA*Δ*hpf*Δ*rmf*Δ*rna*)^5^ and incubated with RNase I. RNase I activity was measured as described previously^28^. RNase I activity decreased in the presence of a higher proportion of hibernating ribosomes (Fig. 1c), indicating that they inhibit RNase I activity.

### RNase I does not interact with translating ribosomes

Our *in vivo* findings showed that hibernating ribosomes interact with RNase I (Fig. 1c and Extended Data Fig. 2). To clarify whether this association is caused by the direct interaction or is mediated indirectly by cellular co-factors, we reconstituted *in vitro* hibernating ribosomes by adding RMF and HPF to 70S ribosomes (Supplementary Fig. 1b) and prepared a catalytically inactive mutant of RNase I (H55F/H133F) (hereafter, ‘RNase I mutant’). The RNase I mutant bound to hibernating ribosomes in the absence of additional factors, indicating that RNase I directly interacts with the ribosome (Fig. 2a). In addition, the RNase I mutant bound almost equally well to both the 70S and hibernating ribosomes. This result aligns with previous reports showing that RNase I associates with ribosomes^26,27^, and suggests that ribosome dimerization into the 100S form is not required for RNase I binding. Moreover, it raised the possibility that RNase I binding depends on the translational state of the ribosome rather than on its dimerization. RNase I bound less efficiently to 70S ribosomes isolated from exponentially growing cells, where most ribosomes are actively translating^33^, than to ribosomes from hibernating state (Extended Data Fig. 2). In the binding assay of RNase I in the presence or absence of *gfp* mRNA using an *in vitro* translation system (PURE system) (Fig. 2b), RNase I binding was markedly reduced when mRNA was added (Fig. 2c). A clear inverse correlation was observed between mRNA concentration and RNase I binding: the higher the mRNA concentration, the lower the level of RNase I binding, indicating that active translation interferes with RNase I binding to ribosomes (Fig. 2d). In addition to the previous studies^26,27^, our experimental design enabled a direct comparison between translationally active and inactive states and revealed that RNase I does not readily bind to ribosomes during the translation process. Furthermore, this binding was restored upon addition of RMF and HPF to ribosomes in the PURE system, which suppressed translation (Fig. 2e), supporting our *in vivo* observation that RNase I interacts more strongly with hibernating ribosomes than with ribosomes in the exponential phase (Fig. 1c and Extended Data Fig. 2). Together, these results provide a mechanistic link between RNase I suppression by hibernating ribosomes and the enhanced biofilm-forming phenotype (Fig. 1a, b).

**Fig. 2.**
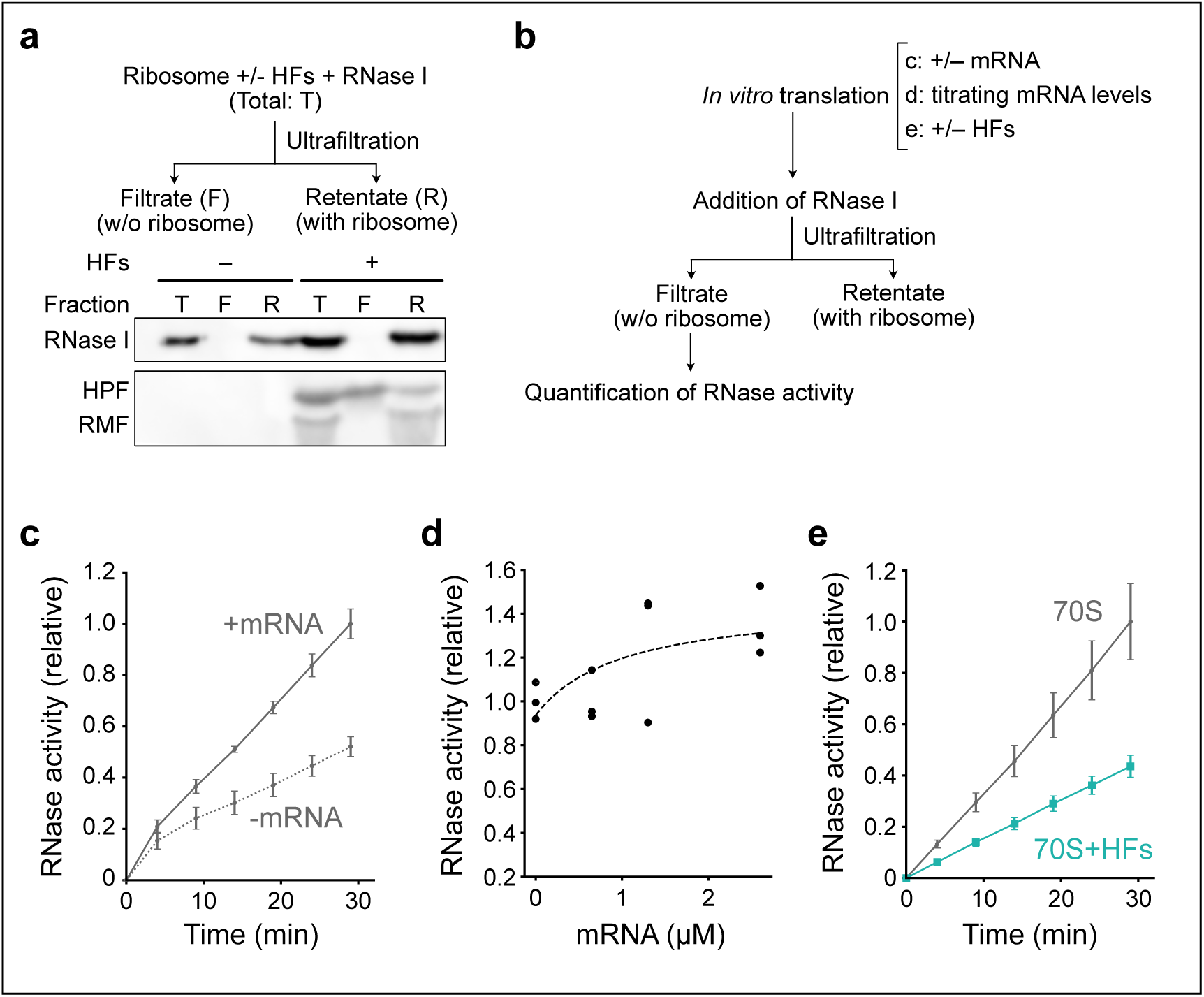
RNase I does not stably associate with translating 70S ribosomes. **a**, Binding ability of 70S ribosomes and *in vitro* reconstituted hibernating ribosomes to RNase I. ‘T,’ ‘F,’ and ‘R’ denote the total mixture, filtrate fraction, and retentate fraction, respectively. **b**, Schematic of the *in vitro* assay used to measure RNase I binding to ribosomes in the PURE system, corresponding to the experiments shown in **c-e**. **c**, RNase I binding with 70S ribosomes in the presence or absence of mRNA. **d**, RNase I binding with increasing mRNA concentrations. **e**, RNase I binding in the presence or absence of RMF and HPF (HFs).

### Structure of the hibernating ribosome bound to RNase I

To elucidate how RNase I interacts with hibernating ribosomes, we reconstituted a complex of the dimerized *E. coli* 100S ribosome and the RNase I mutant. The purified RNase I mutant was mixed with 100S hibernating ribosomes isolated from *E. coli* W3110 cells in the stationary phase. The mixture was then applied to the EM-grids for cryo-EM observations. We collected cryo-EM movies and performed single-particle analysis of the 100S•RNase I complex using two- and three-dimensional classifications (Extended Data Fig. 3). Since the 100S ribosome easily dissociates into two 70S ribosome monomers at low concentrations and tends to exhibit a preferred orientation on EM-grids^32,34^, we extracted 70S ribosome particles from the micrographs. The density map of the 70S•RNase I complex, in which both hibernation factors HPF and RMF were clearly visible, was selected and refined. The resulting complex was obtained from 351,633 particles, yielding a density map at a resolution of 2.4 Å. However, because the density of RMF was not fully occupied in this complex, the 3D classification was repeated to enrich the density, finally yielding a refined map to a resolution of 2.7 Å consisting of 50,955 particles (Fig. 3a, Extended Data Table 1, and Extended Data Figs. 3 and 4a, b).

**Fig. 3.**
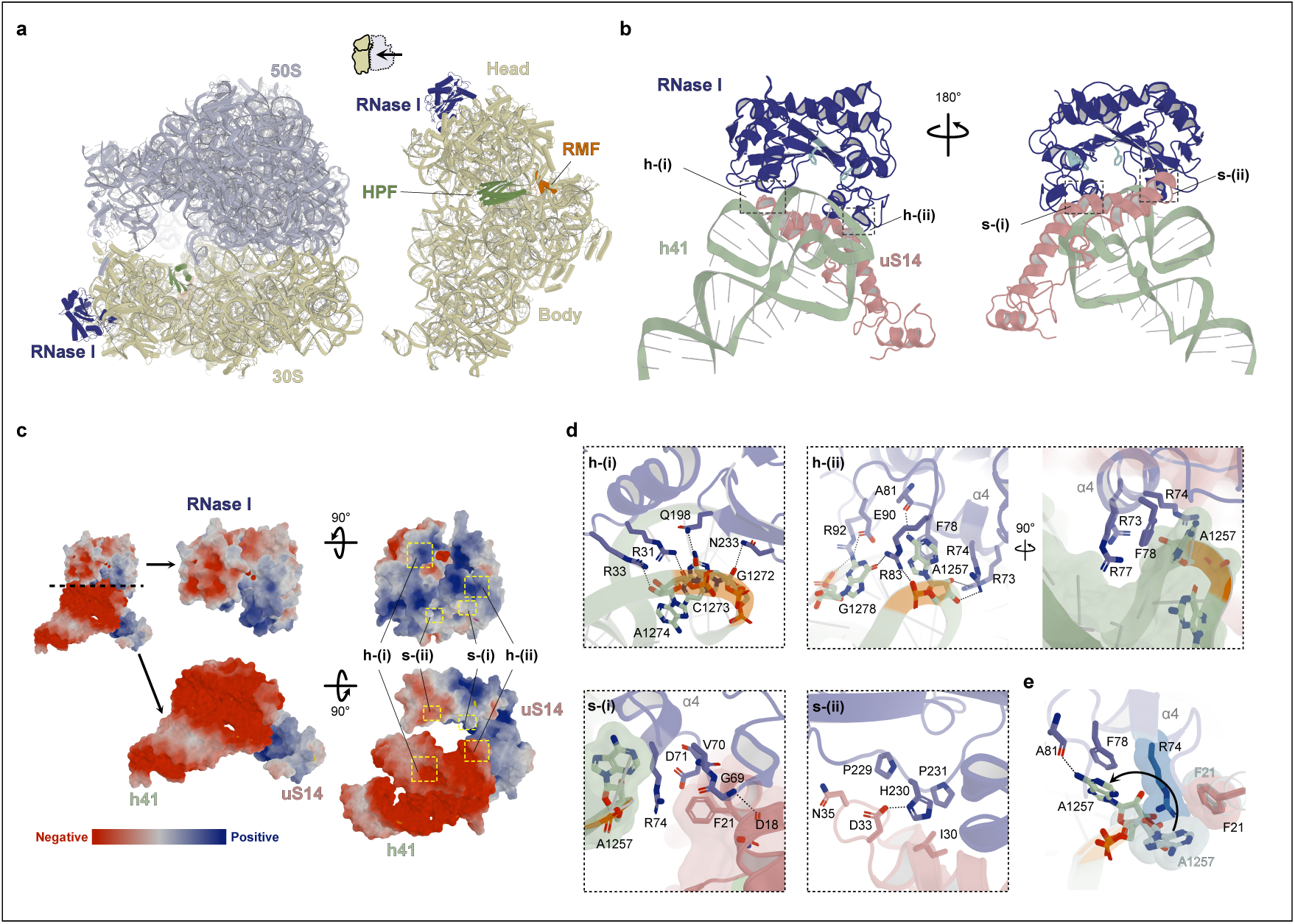
Structure of the RNase I-bound hibernating ribosome complex. **a**, Overall view of the RNase I-bound hibernating ribosome, showing the structure of the *E. coli* 70S ribosome (50S: light blue, 30S: light yellow) complexed with RNase I (dark blue), HPF (green), and RMF (orange) (left). A view of **a** from a different angle with the 50S subunit omitted for clarity (right). **b**, Close-up views of the interaction between RNase I and h41 (light green) of the 16S rRNA or the ribosomal protein uS14 (light pink) from two different angles. The active centre of RNase I is represented by stick models with pale blue. The h41 binding site is shown in h-(i) and h-(ii), and the uS14 binding site is shown in s-(i) and s-(ii). **c**, The electrostatic potential surface is shown in the same direction as **b**. Positive, negative, and neutral potentials are shown in blue, red, and white, respectively. Each binding site is illustrated by a yellow-dashed rectangle. **d**, Detailed views of the binding sites of RNase I on h41 and uS14. The amino acids and nucleotides involved in these bindings are represented by stick models, and the N, O, and P-atom are coloured blue, red, and orange, respectively. For easier understanding of the interactions, transparent surfaces are overlaid on the ribbon and stick models (h-(ii), s-(i)). The hydrogen bonds are indicated using dashed lines. **e**, Structural comparison of A1257 of h41 and F21 of uS14 between our model and a 70S ribosome (PDB 8SYL). The 8SYL ribosome is coloured whitish blue. Transparent surfaces are overlaid on stick models of the base of A1257 (8SYL), side chains of R74 (RNase I) and F21 (our model and 8SYL). The conformational change in A1257 is indicated using a curved arrow.

The structure of the RNase I mutant-bound hibernating complex revealed that the RNase I mutant was bound to the head domain of the 30S subunit (Fig. 3a and Extended Data Fig. 4c). As well as the structures of *E. coli* hibernation factors-bound ribosome complex, the factors HPF and RMF interact with the mRNA channel between the 30S head and body domains, and with the anti-Shine Dalgarno (anti-SD) region between the 30S head and the platform binding centre (PBC), respectively^32,34,35^. Structural characterization further revealed that the C-terminal residues R69 and Y71 of the ribosomal protein bS21 interacted with residues C1100 and A1167 of the 16S rRNA near the PBC (Extended Data Fig. 5a). Compared with the previous structure (PDB 6H4N^32^), our structure shows subtle differences in the 30S subunit, which rotates clockwise by ∼1°, while the head domain swivels by ∼2° (Supplementary Fig. 2). The slight conformational changes caused by the RNase I mutant binding may contribute to the structural arrangement of the PBC, promoting the stability of bS21 (Extended Data Fig. 5a). In addition, the HPF stabilized the inter-subunit bridge B2a by changing the conformation of the decoding centre and bridging to the 50S subunit (Extended Data Fig. 5b). Since the binding site of RNase I is located on the opposite side of the binding regions for HPF and RMF, RNase I and the hibernation factors did not interfere with each other (Fig. 3a). In addition, RNase I could be accommodated in the previous 100S ribosome (PDB 6H58^32^) without steric clashes.

The overall shape of the RNase I mutant determined in this study is nearly identical to the previous crystal structure (PDB 2Z70^36^). *E. coli* RNase I has a characteristic arch-like architecture with active residues H55 and H133 and two piers. In our complex structure, the piers of RNase I interact with the ribosome, and the active centre remains over 6.5 Å from the ribosome, allowing 16S rRNA to evade cleavage (Fig. 3b). In addition to helix 41 (h41) of 16S rRNA, as indicated in a previous report^28^, the ribosomal protein uS14 interacts with the piers of the arch. Analysis of the binding interfaces and areas using PDBe-PISA^37^ and APBS^38^ suggested that RNase I binds primarily to h41 and secondarily to uS14, through shape complementarity (Fig. 3c, Extended Data Fig. 6 and Extended Data Table 2).

### Interaction between the hibernating ribosome and RNase I

The following details are provided regarding the structural basis for the interaction of h41 and uS14 with RNase I. The h41 binds to RNase I by bridging the arch piers (Fig. 3b and Extended Data Fig. 6a, b). The polar amino acid residues R31, R33, Q198, and N233 of RNase I form hydrogen bonds with nucleotides G1272–A1274 in the h41 stem region. Although these amino acids do not directly contact with the groove of the h41 stem loop, they exploit the arch-shaped features of RNase I to interact with only one strand of h41 (Fig. 3c and 3d [h-(i)]). In the other binding region, RNase I interacts with A1257 and G1278 at the kink in the h41 stem. These two nucleotides are unpaired, interacting with and stabilizing the binding of RNase I. In particular, the residue R83 forms hydrogen bonds with both A1257 and G1278 and exhibits stacking interactions with the base of A1257 (Fig. 3c and 3d [h-(ii)]). Interestingly, upon RNase I binding, the A1257 rotates approximately 100° towards the h41 stem-loop to avoid collision with R74 (Fig. 3e), indicating that the ribosome adapts its structure to accommodate RNase I by partially altering its native interactions^39^. This rearrangement creates a pseudo-major groove comprising A1257–G1260 and A1274–G1278, into which helix α4 of RNase I fits via shape complementarity (Fig. 3d [h-(ii)] and Extended Data Fig. 7). In this structure, F78 forms a π-stacking interaction with the base of A1257 on the opposite side of R83, while R73 and R74 from helix α4 form hydrogen bonds with the ribose of A1257 (Fig. 3d [h-(ii)]). The conformational change of A1257 induced by RNase I binding disrupts its native interaction with and F21 of uS14 (Fig. 3e), creating a cleft between h41 and uS14 is filled by R74 of RNase I (Fig. 3d [s-(i)]). Instead of binding to A1257, R74 of helix α4 binds to F21. Additionally, the loop containing G69–D71 of RNase I contacts residues D18–A22 in the bent region of the N-terminal helix of uS14 (Fig. 3b and 3d [s-(i)]). According to PDBe-PISA analysis, the N-terminal helix of uS14 also binds to the RNase I by bridging both arch piers (Extended Data Fig. 6a, c). Residues P229–P231 in the proline/glycine-rich long loop between β5 and β6 of RNase I form electrostatic interactions with residues I30–N35 of uS14, including a hydrogen bond between H230 and D33 (Fig. 3c, 3d [s-(ii)]). Together, these findings provide structural insights into how hibernating ribosomes associate with RNase I in a novel fashion that suppresses its catalytic activity, thereby protecting themselves from degradation.

### RNase I targets dissociated ribosomal subunits

Although RNase I is inhibited on hibernating ribosomes, how RNase I targets and degrades rRNA remains unclear. The association of ribosomal subunits provides protection against RNases, as rRNA is shielded by ribosomal proteins^40^. Consistent with this, we observed that RNase I hardly degrades rRNA within 70S ribosomes *in vitro* (Extended Data Fig. 8a). Since ribosomes are estimated to outnumber RNase I by approximately 100-fold in *E. coli* cells^28^, the observation that a three-fold excess of RNase I is required to degrade 70S ribosomes *in vitro* suggests that such degradation is unlikely to occur under physiological conditions (Extended Data Fig. 8a). This implies that RNase I preferentially cleaves rRNA within dissociated 30S and 50S subunits, rather than intact 70S ribosomes (Fig. 4a). Several rRNA fragments were observed, suggesting the presence of specific RNase I cleavage sites at subunit interfaces. Notably, this degradation occurred at equimolar concentrations of RNase I and ribosomes, in contrast to the degradation of 70S ribosomes (Fig. 4a and Extended Data Fig. 8a). Although dissociated subunits are relatively scarce in exponentially growing cells, they transiently appear and are highly susceptible to RNase I-mediated degradation, making them likely targets before assembly into 70S ribosomes. Furthermore, as mentioned above, hibernating ribosomes are less prone to dissociate into subunits. This feature aligns with the observed decrease in 2′,3′-cNMP levels during the stationary phase^29^, when hibernating ribosomes accumulate. To confirm *in vivo* that subunit dissociation alters the susceptibility of rRNA to RNase I cleavage, we treated *E. coli* cells with puromycin, which facilitates the release of nascent polypeptides and promotes ribosome dissociation into subunits^41^. We then extracted 2′,3′-cNMPs from the cells and analysed them by liquid chromatography–mass spectrometry (LC–MS). Accumulation of dissociated subunits by the addition of puromycin led to increased 2′,3′-cAMP and 2′,3′-cGMP levels in RNase I-expressing cells, indicating enhanced rRNA degradation following subunit dissociation (Fig. 4b). Although 2′,3′-cCMP and 2′,3′-cUMP levels also increased in puromycin-treated RNase I-expressing cells, their levels were lower (Extended Data Fig. 8b). To summarize the above, RNase I degrades rRNA within dissociated subunits and generates 2′,3′-cNMPs in the exponential phase, contributing to the suppression of biofilm formation. We also detected 2′,3′-cNMPs in puromycin-untreated cells, albeit at lower levels. This is consistent with a previous study showing the presence of 2′,3′-cNMPs during the exponential phase^29^.

**Fig. 4.**
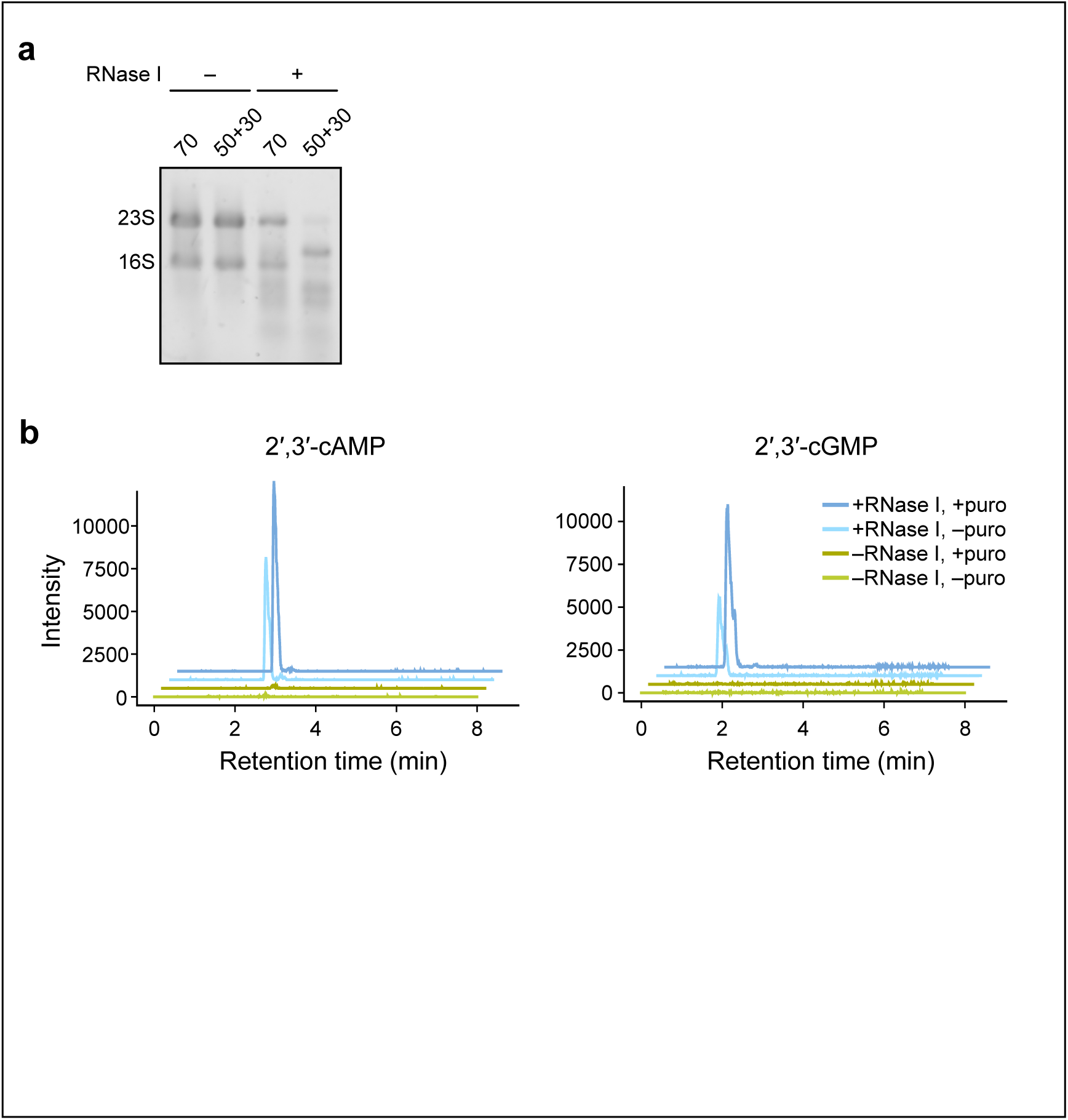
RNase I degrades rRNA from dissociated subunits. **a**, Comparison of rRNA cleavage efficiency of RNase I between intact 70S ribosomes and dissociated 50S and 30S subunits. ‘70’ indicates rRNAs from intact 70S ribosomes, whereas ‘50+30’ indicates rRNAs from dissociated subunits. **b**, RNase I-mediated RNA cleavage in cells upon ribosome dissociation. Δ*rna* cells carrying either an empty vector or a plasmid expressing RNase I were treated with or without puromycin (puro), and the peaks of 2′,3′-cAMP and 2′,3′-cGMP in cell extracts were detected by LC–MS. Extracted ion chromatograms for 2′,3′-cAMP (upper; *m/z* 330.0598 [M+H]^+^) and for 2′,3′-cGMP (lower; *m/z* 346.0547 [M+H]^+^) are shown.

## Discussion

Our study demonstrated that hibernating ribosomes suppress RNase I activity by sequestering it in the head domain of the 30S subunit, thereby inhibiting rRNA cleavage and promoting biofilm formation. These findings revealed that RNase I activity is passively regulated by the ribosomal state; it is not accessible to actively translating ribosomes, and is sequestered by hibernating ones. In addition, this study provides mechanistic and structural insights into how RNase I binds to the 30S subunit of the hibernating ribosome^26,27^. Although the ribosome protects itself from RNase I-mediated cleavage, it is eventually degraded^31^, consistent with our observation that RNase I gradually degrades rRNA in hibernating ribosomes during prolonged stress (Extended Data Fig. 1). Dimeric 100S ribosomes progressively dissociate during prolonged stress, finally exposing their subunits^31^. Our study highlights the potential physiological role of RNase I in binding to hibernating ribosomes. RNase I may not only lose catalytic activity upon binding to hibernating ribosomes but also remain nearby, poised for rapid degradation of ribosomes when required. This is consistent with a previous report suggesting that rRNA degradation contributes to the recycling of phosphates and nucleotides during starvation^42^. Furthermore, RNase I cleaves of rRNAs within dissociated ribosomal subunits into fragments (Fig. 4a), which are then likely processed by exoribonucleases such as RNase R or PNPase^40^, and/or by RNase I itself, thereby contributing to the clearance of excess ribosomes and adaptation to prolonged stress. This degradation may also potentially serve as a programmed response facilitating the transition to cell death when recovery is not possible. Conversely, upon recovery from stress, the increase in 2′,3′-cNMP levels through ribosome degradation may promote biofilm disassembly, allowing cells to prepare for regrowth.

Ribosome dissociation occurs naturally during the translation cycle via the ribosome recycling factor, and accidentally stalled ribosomes on mRNA, on the other hand, are rescued and split by ribosome rescue systems^43–45^. The conserved GTPase HflX also functions as a ribosome-splitting factor for mRNA-vacant ribosomes^46^. Given that RNase I efficiently cleaves rRNA in dissociated subunits (Fig. 4a, b) and the estimated excess of ribosomes over RNase I molecules within the cell^28^, targeting these fleeting subunits rather than stable 70S complexes provides an efficient surveillance mechanism. This model could directly link ribosomal conformational dynamics and quality-control processes to rRNA degradation.

One of the most remarkable findings of this study is the elucidation of how RNase I binds to the hibernating ribosome. Our structure reveals that the RNase I-binding site lies within the 30S head domain, the conformation of which constantly fluctuates during translation elongation because of swivel and rotation^47–49^. Such dynamic fluctuations likely destabilize of the head domain during active translation^47^, thereby precluding its interaction with RNase I (Fig. 2c, d). In contrast, when ribosomes are not actively engaged in translation or when ribosomes enter the hibernation state, the fluctuation of the head domain is suppressed (Fig. 5 (i) and (ii)), facilitating stable RNase I association. A molecular dynamics simulation of the translation initiation complex predicted conformational fluctuations in the C1254–C1284 region of h41^49^. Notably, this region includes A1257–G1260 and A1274–G1278, which interact directly with RNase I in our structure (Fig. 3), suggesting that RNase I binding stabilizes these otherwise dynamic regions of h41. This structural information raises the possibility that RNase I binding suppresses the dynamics of the head domain and finally inhibits translation (Fig. 5(iv)). Consistent with this, RNase I inhibited protein synthesis *in vitro* in a concentration-dependent manner (Extended Data Fig. 9). As the concentration of the RNase I mutant increases, the binding–dissociation equilibrium may shift towards the bound state, allowing RNase I to engage ribosomes in transiently non-translating states or promote inactivation. Together, our structural and biochemical data highlight a bidirectional relationship between ribosomal conformational states and RNase I activity.

**Fig. 5.**
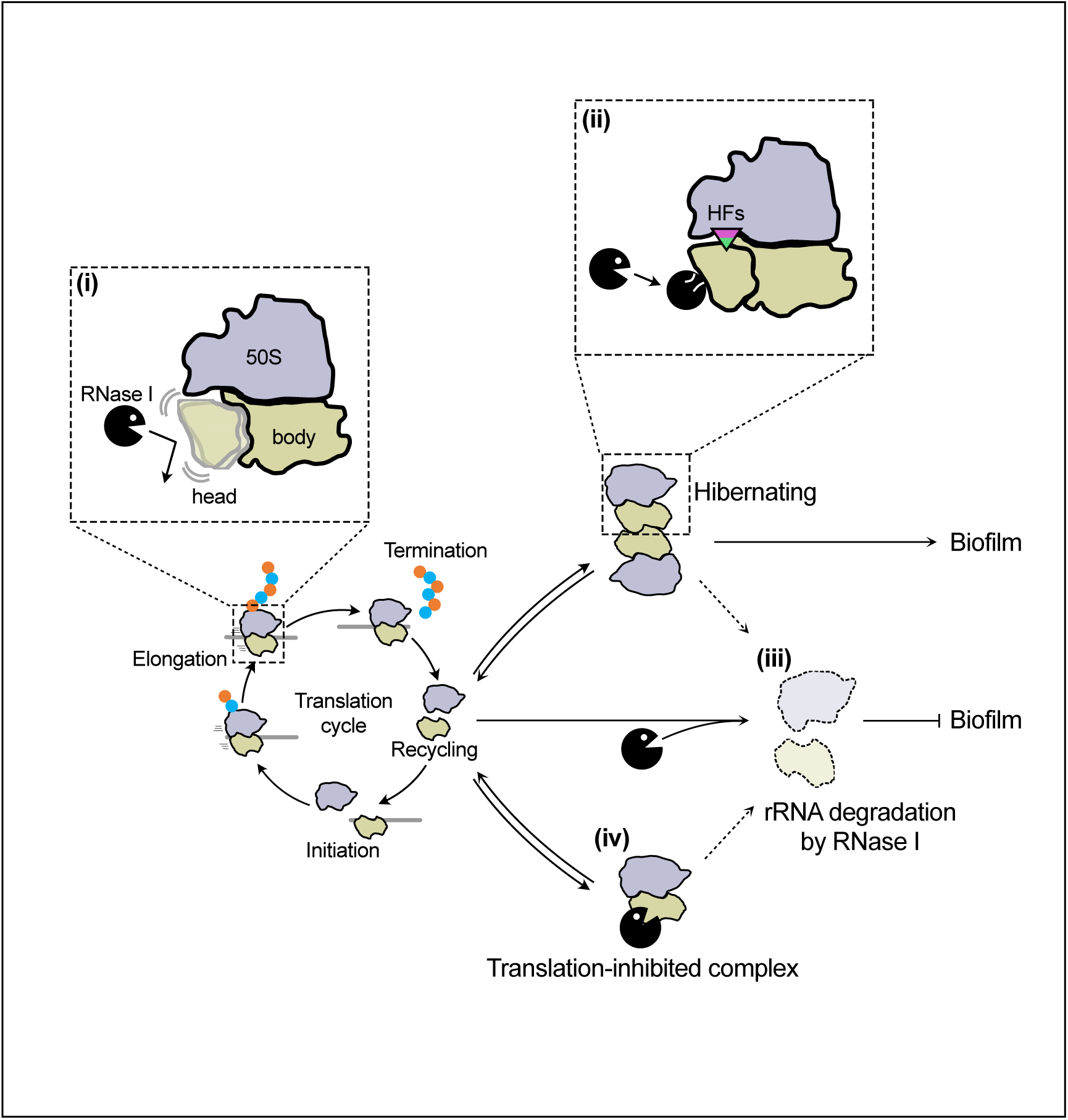
Regulation of RNase I by ribosomal states. Schematic of the interaction between RNase I and ribosomes. (i) In the translating ribosome, the 30S head swivels relative to the body, rendering the RNase I binding site unstable and precluding productive engagement (related to Fig. 2b– d). (ii) Binding of HFs locks the head domain in a rigid conformation in the 100S complex, stabilizing RNase I binding (related to Figs. 1c, 2e, 3). (iii) Dissociated 30S and 50S subunits are recognized by RNase I and rRNAs are degraded (related to Fig. 4). (iv) The binding of RNase I to the non-translating 70S ribosome results in a translation-inhibited complex (related to Extended Data Fig. 9). Such complexes, as well as the hibernating ribosomes, may subsequently become targets for RNase I-mediated rRNA degradation if they dissociate into subunits.

Our cryo-EM structure shows that RNase I directly interacts with the ribosomal protein uS14, providing new insight into the evolutionary plasticity of this conserved ribosomal component. Although the gene encoding uS14 is essential^50^, its precise role in translation remains poorly understood. uS14 proteins are categorized into three classes based on the presence or absence of a zinc-binding motif: C+, C– short, and C– long^51^. Compared with *Thermus thermophilus* uS14 (C+ class) in the crystal structure of the 70S ribosome (PDB 4V8H^34^), *E. coli* uS14 (C– long class) possesses a distinct domain insertion that forms the RNase I binding interface (Extended Data Fig. 10a). No RNase I homologue was identified in *T. thermophilus*, suggesting that the interaction between RNase I and uS14 may have evolved specifically in certain bacterial lineages. Additionally, the presence of C+/– uS14 paralogue pairs in some bacteria, along with reported differences in their functional properties^51–53^, further supports the idea that uS14 is structurally adaptable, potentially enabling diverse functional acquisitions, including, in *E. coli*, facilitating interaction with RNase I.

The binding interface between uS14 and RNase I raises the question of whether the interaction depends solely on ribosomal features or also on structural adaptations in RNase I. Structural comparison of *E. coli* RNase I and human RNase T2 revealed that *E. coli* RNase I contains an internal insertion region (Extended Data Fig. 10b). This insertion (S64–F78) forms a small domain of two short helices and connecting loops, which contributes to the binding interface with both uS14 and h41 (Fig. 3d [s-(i)] and [h-(ii)], Extended Data Figs. 6, 10b), suggesting that the ribosome-binding specificity of RNase I is an evolutionary acquisition, likely shaped by coevolution with ribosomal elements such as uS14 and h41. Although the overall structures of ribosomes are widely regarded as highly conserved across species, local variations in rRNA and ribosomal protein architecture have been known. Our findings suggest that ribosomes may have adapted their structures not only for translation but also for interacting with regulatory factors like RNase I.

Recently, cryo-EM structures of *Bacillus subtilis* RNase R in complex with the 30S subunit were determined, showing that it targets the anti-Shine-Dalgarno (anti-SD) sequence and helix 44 within the decoding centre for rRNA degradation^54^. The RNase R binding site overlaps with that of RMF (Extended Data Fig. 10c), indicating that RMF functions as an inhibitor that protects hibernating ribosomes from RNase R-mediated degradation^5^. Our structure represents an inactive state in which RNase I binds to the 30S head domain in a manner that suppresses its catalytic activity. This interaction may allow RNase I to remain poised for activation upon ribosome dissociation.

In summary, our findings provide insights into the mechanisms by which ribosomes inhibit RNase I activity to preserve ribosome integrity under stress. This study highlights ribosome–nuclease crosstalk, offering a novel perspective on how ribosomes not only execute translation but also regulate RNA metabolism in response to environmental stimuli.

## Methods

### Construction of strains and plasmids

The strains and plasmids used in this study are listed in Supplementary Table S1. RNase I, RaiA, RMF, or HPF-deficient strains (Δ*rna*, Δ*raiA*, Δ*rmf*, or Δ*hpf*, respectively) were obtained from the Keio Collection, provided by the National BioResource Project-E. coli (National Institute of Genetics, Japan)^55^. The kanamycin-resistance gene cassette of the Keio Collection strains was removed by homologous recombination using plasmid 707-FLPe (Gene Bridges). Double (Δ*raiA*Δ*rna*) and quadruple knock-out strains (Δ*raiA*Δ*hpf*Δ*rmf*Δ*rna*) were constructed by homologous recombination using the Red/ET system (Gene Bridges).

The plasmid pET-*gfp* was constructed by inserting the *gfp* gene between the NcoI and NotI sites of pET24d(+). The open reading frame (ORF) and promoter region of the *rna* gene with a C-terminal FLAG tag were PCR-amplified and inserted into the pUC18 vector using the seamless cloning method (In-Fusion HD Cloning Kit, TaKaRa Bio). For expression of His-tagged RNase I, RMF, and HPF, the respective ORFs were PCR-amplified and inserted between the NcoI and EcoRI sites of pET24d(+). A plasmid expressing a catalytically inactive RNase I mutant was constructed by substituting the codons for H55 and H133 with phenylalanine in the *rna* gene on pET-*rna*.

### Preparation of RNase I and hibernation factors

*E. coli* BLR(DE3) strains transformed with plasmids expressing either His-tagged RNase I, RMF, or HPF, were cultured in L-broth supplemented with 30 µg/mL kanamycin. When an optical density at 660 nm (OD_660_) reached 0.5, isopropyl β-D-1-thiogalactopyranoside (IPTG) was added to the culture to a final concentration of 1 mM, and the cells were further cultivated for 3 h at 37 °C. RNase I was purified as described previously^16^. For the purification of RMF and HPF, buffer conditions were followed as described in a previous publication^34^. For the purification of RMF, an equal volume of 2 M LiCl was added to the supernatant after sonication and stirred overnight at 4 °C, followed by purification^56^.

### Isolation of ribosomes and sucrose density gradient centrifugation

*E. coli* 70S ribosomes were prepared from the Δ*raiA*Δ*hpf*Δ*rmf*Δ*rna* strain as described^57^. *E. coli* 100S ribosomes for cryo-EM were prepared as follows: 2 mL of an overnight preculture of *E. coli* K-12 W3110 strain was obtained using medium E (containing MgSO_4_, citric acid, K_2_HPO_4_, and NaNH_4_HPO_4_) supplemented with 2% polypeptone and 0.5% glucose^58^. Mass cultures were performed in 100 mL of the same medium at 37 °C for 24 h. Cells were harvested and suspended in an association buffer (20 mM Tris-HCl (pH 7.6), 15 mM Mg(OAc)_2_, 100 mM NH_4_OAc, and 6 mM β-mercaptoethanol). Cells were vortexed with an approximately equal volume of glass beads (212–300 µm, Merck KGaA) for 3 min. The homogenate was centrifuged at 12,000 × *g* for 8 min at 4 °C. The supernatant was applied on top of a 5–20% linear sucrose density gradient in the association buffer and centrifuged using an SW 41 Ti rotor (Beckman Coulter, Inc.) at 285,000 × *g* for 1.5 h at 4 °C. After centrifugation, the absorbance was measured at 260 nm using a UV-1800 spectrophotometer (Shimadzu Co.) to obtain ribosome profiles. The 100S ribosomal fractions were obtained based on the profiles. The solution containing 100S ribosomes was concentrated, and sucrose was removed using a centrifugal filter Amicon-100K (Merck KGaA).

### Extraction of total RNA from E. coli cells

*E. coli* strains were cultured in L-broth at 37 °C until an OD_660_ reached 0.5. Cells were washed, suspended in M9 minimal medium (supplemented with 0.4% glucose and 0.2% casamino acids), and further incubated for 0, 2, 4, or 24 h. The cells were harvested at the indicated time points, and total RNA was extracted using an RNeasy Mini Kit (QIAGEN). Total RNA was analysed by electrophoresis on a denaturing 1% agarose gel containing 2% formaldehyde in MOPS buffer or an Agilent 2100 Bioanalyzer (Agilent Technologies).

### Quantification of biofilm formation

Biofilm formation was quantified by crystal violet staining following a published procedure^59^ with slight modifications as described^60^. *E. coli* was cultured in 10 mL of L-broth at 37 °C to an OD_660_ of approximately 0.5. The culture was then diluted 1:100 in M9 minimal medium with 0.2% casamino acids, and 100 µL aliquots were dispensed into a U-bottom 96-well plate (FALCON #351177, Corning). The plates were incubated statically at 30 °C for 24 h. Following incubation, non-adherent cells were decanted by submerging the plates twice in a water bath. To stain the biofilm, 125 µL of 0.1% crystal violet was added to each well and incubated at room temperature for 15 min on a rocking shaker. Excess stain was removed by washing the plates twice in a water bath. The plates were air-dried overnight. To solubilize the remaining crystal violet, 125 µL of 30% acetic acid was added to each well, and the plate was shaken for 15 min. Absorbance was measured at 570 nm using a microplate reader (Infinite F200, TECAN).

### In vitro RNA degradation and RNase I binding assays

RNA degradation assays were performed based on a previous study^28^. For the inhibition assay (Fig. 1c), 0.075 pmol RNase I was incubated with 0.69 pmol of ribosomes. Ribosomes were prepared from Δ*raiA*Δ*rna* or Δ*raiA*Δ*hpf*Δ*rmf*Δ*rna* strains following a published method^31^. Mixtures were incubated at 30 °C for 30 min, and the activity of free RNase I was quantified using the RNase Activity Assay Kit (Abcam).

For the binding assay (Extended Data Fig. 2), spheroplasts of *E. coli* cells were prepared as follows: cells were washed with 10 mM Tris-HCl (pH 8.0), and the pellet was suspended in 500 µL of the buffer containing 10 mM Tris-HCl (pH 8.0) and 0.5 M sucrose. After 30 min of incubation, 500 µL of the buffer containing 10 mM Tris-HCl (pH 8.0) and 2 mM EDTA was added, further incubated for 15 min, and centrifuged at 5,000 rpm for 10 min. The pellet was suspended in 500 µL of water and centrifuged at 5,000 rpm for 10 min. The lysate was loaded onto an ultrafiltration device (300,000 MWCO; PALL) and centrifuged at 14,000 × *g*. After centrifugation, the retentate and filtrate were separately collected. The amount of RNase I bound to ribosomes was detected by western blotting or the RNase Activity Assay Kit.

The binding assay (Fig. 2a) was performed as follows: 20 µM of 70S ribosomes were mixed with purified RMF and HPF (at least 10-fold molar excess over 70S ribosomes) to reconstitute hibernating ribosomes *in vitro*. Thereafter, 20 µM of 70S ribosomes or the reconstituted hibernating ribosomes were mixed with 120 µM of RNase I mutant and incubated at 37 °C for 30 min. The mixture was then ultrafiltered at 14,000 × *g*, and the retentate and filtrate were collected separately. The RNase I mutant and ribosomal protein (RpsB) in each sample were detected using western blotting. To detect His-tagged RNase I, anti-6x histidine MoAb (9C11) (FUJIFILM Wako Pure Chemical Corporation) and anti-mouse IgG, HRP-Linked Whole Ab Sheep (Cytiva) were used as the primary and secondary antibodies, respectively. To detect RpsB, anti-rpsB antibody (Abcam) and anti-rabbit IgG, HRP-Linked Whole Ab Sheep (Cytiva) were used.

The binding assay using *in vitro* translation (Fig. 2b–e) was performed as follows. *In vitro* translation was performed in the presence or absence of *gfp* mRNA using PURE*frex* 1.0 (GeneFrontier), the reconstituted translation system derived from *E. coli*^61^. The *gfp* mRNA was synthesized using the T7 RiboMAX Express Large Scale RNA Production System (Promega), and transcripts were purified using NucleoSpin RNA (MACHEREY-NAGEL). The translation mixture was incubated at 37 °C for 30 min. RMF and HPF were added to ribosomes prior to initiating *in vitro* translation to reconstitute hibernating ribosomes. RNase I mutant was added at a final concentration of 4.8 µM. After 30 min of further incubation, the reaction mixture was ultrafiltered, the filtrate was collected, and the RNase I activity in the filtrate was measured as described above.

For the *in vitro* translation inhibition assay (Extended Data Fig. 9), *in vitro* translation was performed as described above with the following modifications: RNase I mutant and 1 µM of ribosomes were added to the translation mixture and incubated for 30 min at 37 °C. Translation was initiated by adding *gfp* mRNA, and the translation activity was evaluated based on GFP-derived fluorescence.

### In vitro rRNA degradation from split ribosomal subunits

The susceptibility of rRNA in 70S ribosomes, 30S, and 50S subunits to RNase I was assessed as follows: 20 pmol of ribosomes were resuspended in the prokaryotic ribosome dissociation buffer (10 mM Tris-HCl (pH 7.5), 1 mM Mg(OAc)_2_, 100 mM NH_4_Cl, 1 mM DTT, 0.5 mM EDTA) and incubated overnight at 4 °C for 70S ribosome dissociation^62^. Then RNase I (124 pmol) was added and incubated for 30 min at 37 °C. Total rRNA was then extracted using RNeasy Mini Kit, as described above, and analysed by electrophoresis on a denaturing agarose gel.

### In vivo rRNA degradation from split ribosome subunits and detection of 2′,3′-cNMPs using liquid chromatography–mass spectrometry (LC–MS)

Ribosome dissociation *in vivo* was induced by puromycin treatment, based on protocols modified from a previous research^41^. Δ*rna* strains harbouring either the empty vector pUC18 or pUC18-*rna* were cultured in L-broth until an OD_660_ reached 0.5. Puromycin was added to a final concentration of 100 µM, and cells were incubated for 15 min. Cyclic nucleotides were extracted as follows^63^: Cells were centrifuged at 10,000 rpm for 1 min, washed twice with sterile water, and resuspended in 1 mL of 1 M acetic acid. After incubation on ice for 5 min, lysates were centrifuged at 15,000 rpm for 5 min. Supernatants were collected, freeze-dried, resuspended in 100 µL of water, and filtered using a Spin-X Centrifuge Tube Filter (Corning).

Liquid chromatography-electrospray ionization (ESI)-high-resolution mass spectrometry (HRMS) analysis was performed using a SCIEX triple TOF X500R system (SCIEX) equipped with an UFLC Nexera system (Shimadzu). The UFLC system was equipped with a CAPCELLPAK C18 IF column (2.0 mm × 50 mm; Shiseido) and eluted at a flow rate of 0.4 mL/min. The composition of mobile phase A was methanol–water (3:97, v/v) and B was methanol–water (97:3, v/v), each containing 50 mM ammonium acetate and 0.1% acetic acid^64^. The program was a linear gradient of 0–50% B for 5 min, 50–100% B for 1 min, 100% B for 3 min, 100–0% B for 1 min, and 0% B for 5 min. MS and MS/MS parameters were as follows: ESI positive ion mode; acquisition mass range, *m/z* 50–500; TOF accumulation time, 0.1 S; MS/MS accumulation time, 0.1 s; collision energy, −10 eV; ion source gas 1, 60 psi; ion source gas 2, 60 psi; curtain gas 30 psi; ion spray voltage floating, −4500 v; temperature 350 °C. MS data were analysed using an Analyst software (SCIEX).

### Sample preparation and cryo-electron microscopy data acquisition

The ribosome•RNase I complex was reconstructed using 1.2 µM 100S hibernating ribosomes containing HPF and RMF from *E. coli* K-12 W3110 strain and 7.2 µM RNase I in the reaction buffer (20 mM HEPES-KOH (pH 7.6), 10 mM Mg(OAc)_2_, 30 mM NH_4_Cl, and 6 mM β-mercaptoethanol) at 37 °C for 30 min. The complex was diluted by 1.5-fold, applied onto an EM-grid, and plunge-frozen in liquid nitrogen-cooled ethane. Quantifoil R2/2 gold 200 mesh grids (Quantifoil Micro Tools GmbH) glow-discharged for 30 s in a PIB-10 Ion Bombarder (Vacuum Device Inc) were used to apply 2.7 µL sample, pre-incubated for 10 s, blotted for 15 s, and post-incubated for 2 s at 4 °C and 100% humidity before freezing in a Vitrobot Mark IV (Thermo Fisher Scientific). The grids were subsequently clipped into an autogrid cartridge and transferred into a CRYO ARM^TM^ 300 II field emission cryo-electron microscope (JEOL) operating at 300 keV. Data were acquired with a K3 camera (Gatan) using SerialEM software^65^. The cryo-EM movies were acquired at a calibrated magnification of 80,460 and a pixel size of 0.87 Å/pixel to record movies with 70 frames with a total accumulated dose of 50 e^−^/Å^2^/movie.

### Cryo-electron microscopy image processing

All steps of the image processing were performed using cryoSPARC 3.3.1 and 4.0.0 software^66^. 2,000 of 33,302 movies were initially motion-corrected by the patch motion correction job, followed by the patch CTF estimation. A total of 258,744 particles were picked by circular blob (size 260–300 Å) using the blob picker job and extracted at 4-fold decimated pixel size (3.48 Å/pixel). The extracted particles were two-dimensional (2D) classified to remove the classes of 30S subunits and debris. The 70S particles were re-picked at a size of 360 Å using the classes selected as templates, followed by the extraction from micrographs job and 2D classification jobs. Five ab-initio volumes were reconstructed using selected 146,282 particles to obtain the main class average of the 70S ribosome; the classes of 50S subunit and debris were discarded by heterogeneous refinement. After the 70S-like particles were combined using homogeneous refinement, followed by local refinement with signal subtraction on the 30S head, a three-dimensional (3D) classification focused on the head region was performed to obtain RNase I-binding particles. Selected 33,823 particles were extracted at 1.25-fold decimated pixel size (2.175 Å/pixel), the complex could not find the RMF in the volume at a nominal resolution of 2.7 Å. Remaining movies were processed in the same way as described above. Heterogeneous refinement was performed to extract 70S-like classes composed of total 1,501,256 particles from the 2D classes of all 33,294 micrographs. A soft mask focused on the 30S head with hibernation factors and RNase I was generated using UCSF ChimeraX 1.5^67^, before 351,633 particles containing hibernating 70S with RNase I were obtained by the 3D classification, followed by the local refinement with signal subtraction to improve the local resolution of the focused region. Resulting the local 3D volume was refined to a nominal resolution of 2.4 Å, yet the density of RMF remained poor. Two rounds of 3D classification were performed, and particles containing the weak density map of RMF were discarded. 3D classification using the mask on the 30S platform, including S1, further sorted out 50,955 particles, including RMF and part of S1. The final reconstruction was refined to a resolution of 2.7 Å in a homogeneous map and to 2.7 Å in a local map of the mask on 30S head, taking into account the global/local CTF refinements after the particles were re-extracted at 1.4-fold decimated pixel size (1.21 Å per pixel). Since the orientation of both homogeneous and local volumes were slightly different, the local volume was docked into the homogeneous volume using the ‘Fit in Map’ function and resampled to the grid of homogeneous refinement using trilinear interpolation by the ‘volume resample’ operation in ChimeraX 1.6.1^67^.

### Model building and refinement

The structure of the *E. coli* 70S ribosome in this study was built in the 2.7 Å density map obtained from homogeneous refinement in cryoSPARC, using a model of the *E. coli* 70S ribosome structure (PDB 8SYL^5^) as a template. After removing all molecules and ions other than ribosome components of the 8SYL model, the ribosomal subunits, split into 50S and 30S using PyMOL, were docked as rigid bodies into the density map using the ‘Fit in Map’ function in ChimeraX 1.6.1^67^. The model of each subunit was refined with jelly-body restraints using Servalcat 0.4.74^68^. The models were manually corrected for ribosomal proteins according to secondary structure and Ramachandran restraints for better fitting into the density map using the ‘Real Space Refine Zone’ function in Coot 0.9.8.95^69,70^. In addition, the nucleotides of the rRNA chains in the models were manually corrected for syn-or anti-conformations with reference to the differential Fourier map produced by Servalcat. The models of the HPF derived from the crystal structure of *Tth*70S•*Eco*HPF complex (PDB 4V8H^34^) and the RMF derived from the cryo-EM structure of *E. coli* hibernating ribosome complex (PDB 6H4N^70,71^) were rigid-body fitted into the homogenous density map. The crystal structure of *Eco*RNase I (PDB 2Z70^36^) was rigid-body fitted into the EM density map of the local refinement focused on the 30S head region in cryoSPARC; two mutated residues, H55F and H133F, at the active site of the RNase I model were also manually modified and corrected in Coot. These models were merged into the 30S model before it was jelly-body refined by Servalcat, and the 30S binding partners, especially, were manually corrected residue by residue in Coot, together with Ramachandran and secondary structure restraints. After merging the models of the 30S and 50S complexes into the hibernating 70S•RNase I complex, a series of operations in jelly-body refinement and manual correction were repeated several times until the model satisfied its geometry. Magnesium ions and water molecules from 8SYL were docked into the model of the 70S•RNase I complex in Coot. After refining the small molecules merged with the complex using Servalcat, their atomic coordinates were manually corrected or removed to better fit the density map and the differential Fourier map. The resulting model was validated using the comprehensive validation tool for cryo-EM in PHENIX 1.21.1^71,72^ (Extended Data Table S1).

### APBS and PISA calculations

The surface potential values were produced using APBS^38^. The interfacing residues and the buried surface areas were calculated using the protein interface, surfaces, and assemblies (PISA) provided by Protein Data Bank Europe (PDBe)^37^ (https://www.ebi.ac.uk/pdbe/pisa/). Colour coding based on the BSA values was performed on PyMOL using the original Python program colorByBSA.py (https://github.com/minami1009/colorByBSA).

### Preparation of figures

Figures were generated using PyMOL (The PyMOL Molecular Graphics System, Versions 3.0.0 and 3.1.0 Schrödinger, LLC), Chimera X 1.7.1, Matplotlib (Version 3.7.1), and Seaborn (Version 0.13.1) libraries in Python (Version 3.10.12), OmniGraffle (Version 7.18.6), and Affinity Designer (Version 1.10.5).

## Supporting information

Supplementary_bioRxiv_v2

## Data availability

The atomic coordinate of the hibernating 70S ribosome in complex with the RNase I mutant was deposited in the Protein Data Bank Japan (PDBj) under accession code 9IOT. The cryo-EM map was deposited in the Electron Microscopy Data Bank (EMDB) under accession code EMD-60747, and the local map focused on the 30S head was deposited as an additional map.

## Acknowledgments

We thank MG. Gagnon and M. Yao for critical reading of the manuscript and offering suggestions; T. Suzuki, A. Nagao and N. Akiyama for performing the SDGC and isolating of ribosomes; members of the Synthetic Biology Laboratory and T. Shiraishi for supporting and advising on LC-MS analysis; members of the T. Kato group and J. Kishikawa for supporting and advising on the cryo-EM single particle analysis; K. Yamashita for advising on model refinement using Servalcat; and Y. Maki for discussions. We are also grateful to late T. Beppu for the discussion. This work was funded by JSPS KAKENHI (Grant numbers JP20K21266 and JP23H02122) to T.O. and by JST SPRING (Grant number JPMJSP2108) to A.M. The collection of EM movies for single particle analysis was performed using the CRYO ARM 300 II at the JEOL YOKOGUSHI Research Alliance Laboratories of the Graduate School of Frontier Biosciences, The University of Osaka. This research was supported by the Research Support Project for Life Science and Drug Discovery (Basis for Supporting Innovative Drug Discovery and Life Science Research (BINDS)) from AMED under Grant Number JP23ama121001.

## Author contributions

A.M.: conceptualization, data curation, formal analysis, funding acquisition, investigation, methodology, software, visualization, writing -original draft, review and editing. T.T.: data curation, formal analysis, investigation, software, visualization, writing - original draft, review and editing. T.M.: investigation. F.M.: investigation. Z.Y.: methodology. T.F.: methodology, writing - review and editing. To.K.: supervision, writing - review and editing. H.Y.: investigation, resources, writing - review and editing. Ta.K.: funding acquisition, supervision, writing - review and editing. T.O.: conceptualization, data curation, methodology, funding acquisition, project administration, supervision, writing - original draft, review and editing.

## Additional information

**Supplementary Information** is available for this paper.

**Reprints and permissions information** is available at www.nature.com/reprints.

