## Supplementary_bioRxiv_v2 for "Molecular basis of RNase I-mediated rRNA degradation regulated by ribosomes"

**Title**

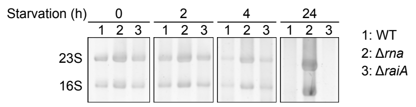

**Extended Data Fig. 1 | rRNA degradation is suppressed in the Δ*raiA* strain under amino acid starvation.** Wild-type, Δ*rna,* and Δ*raiA* strains were cultivated under amino acid starvation condition. Total RNA was prepared at the indicated time points, followed by electrophoresis on a denaturing agarose gel. RNA was visualized using SYBR Gold.

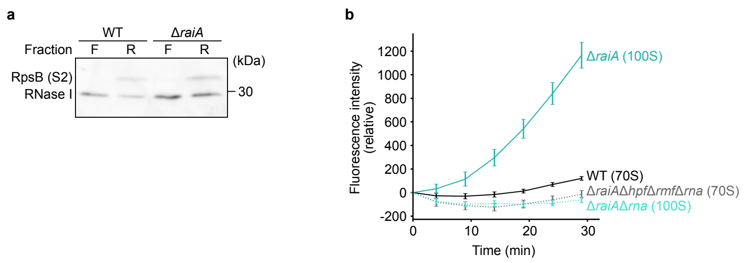

**Extended Data Fig. 2 | RNase I binds exhibits higher affinity for hibernating ribosomes than for 70S ribosomes. a,** Wild-type and Δ*raiA* strains were cultivated in L-broth and amino acid starvation condition, respectively. Then spheroplasts were prepared, lysed, and fractionated by ultrafiltration with a 300 kDa cut-off. RNase I and ribosomes were detected by western blotting using anti-His-tag antibody and anti-ribosomal protein antibody (RpsB), respectively. ‘F’ and ‘R’ indicate the filtrate and retentate fractions, respectively. **b,** Ribosomes were prepared from each strain as indicated, and the amount of RNase I bound to ribosomes was evaluated based on the activity of RNase I (see method “In vitro RNA degradation and RNase I binding assays”).

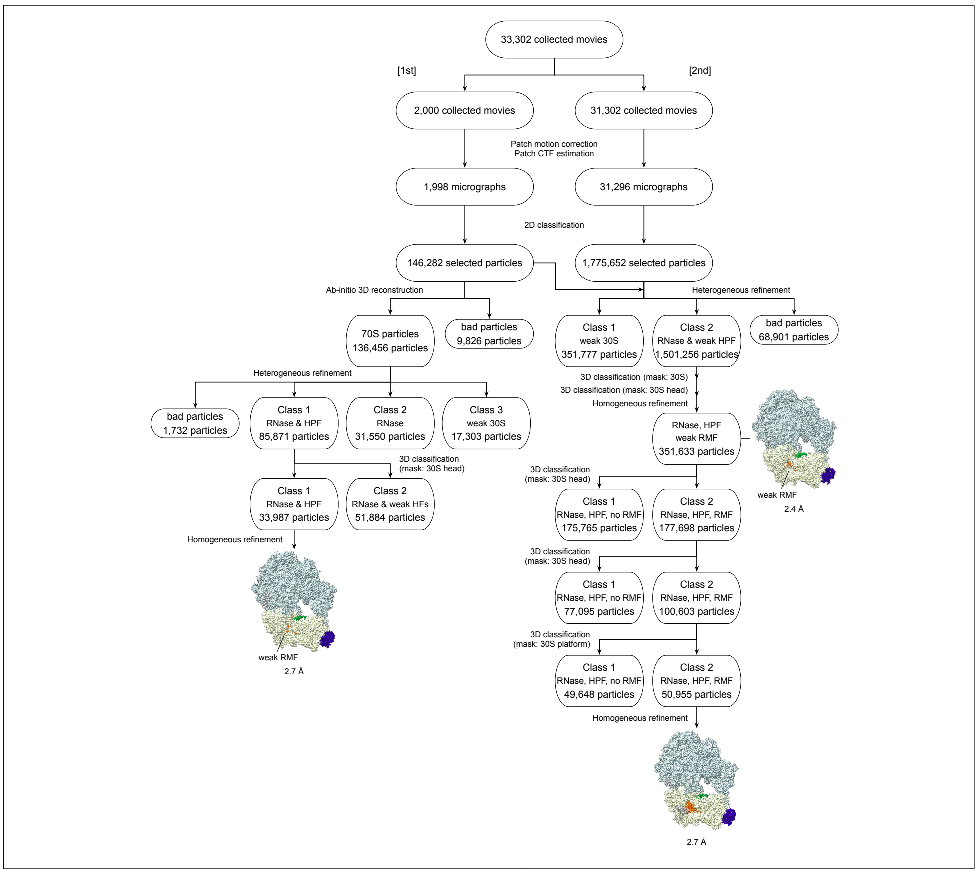

**Extended Data Fig. 3 | Cryo-EM data processing and particle classification workflow**

All processing steps were operated in cryoSPARC v3.3.1 and v4.0.0 according to the flow chart. The numbers of movies, micrographs, and particles are shown in the boxes in each step.

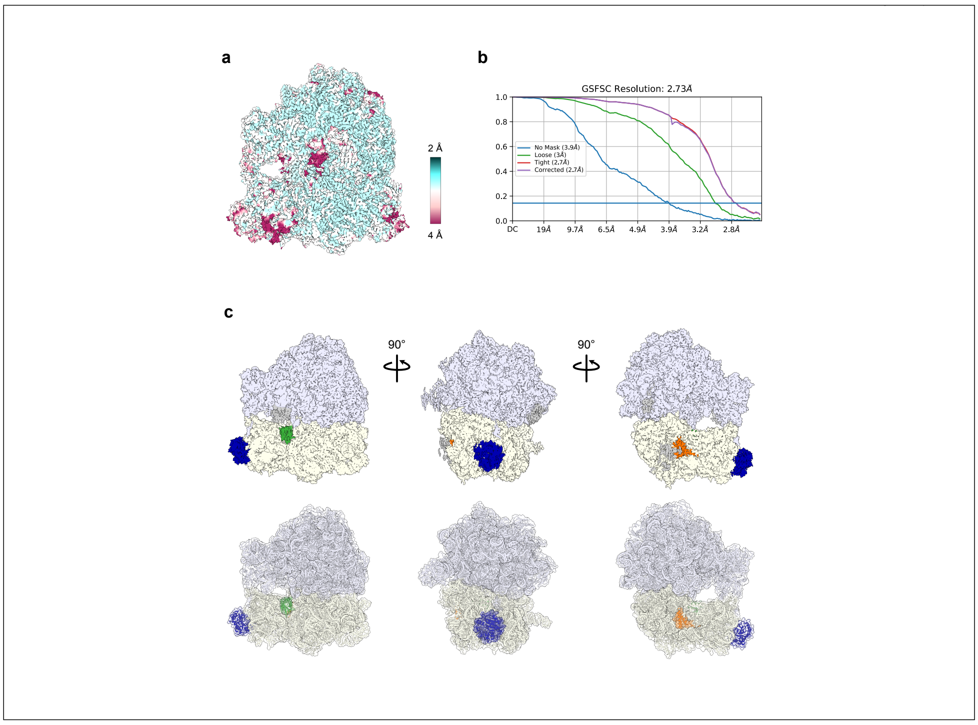

**Extended Data Fig. 4 | Overall view of the volume map and model of the hibernating ribosome•RNase I complex. (related to Fig. 3a)**

**a,** A cryo-EM density map of the hibernating ribosome•RNase I complex. The map is coloured according to the local resolution, as indicated in the colour bar. **b,** Gold-standard Fourier shell correlation (GS-FSC) curves, using GS-FSC = 0.143 for resolution criterion. **c,** From left to right: view from the A-site, front view, and view from the E-site, with volume at the top and model at the bottom. The colour coding is the same as that shown in Fig. 2a.

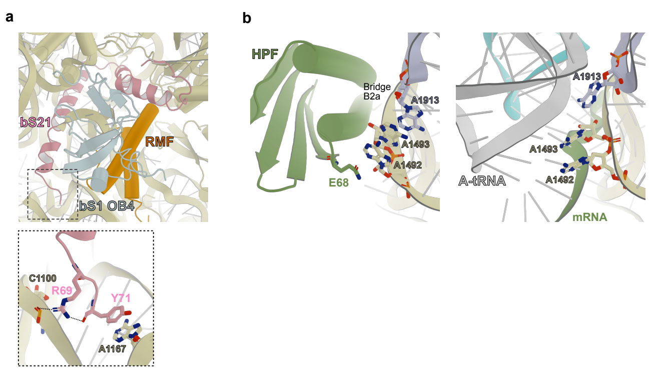

**Extended Data Fig. 5 | Structural characterizations of the hibernating ribosome•RNase I complex.sh**

**a,** Structural organization of RMF (orange), the bS1 OB4 domain (whitish blue), and bS21 (light pink) in the PBC of the hibernating complex. Inset, a close-up view of the interaction between the C-terminal residues R69 and Y71 of bS21 and two nucleotides, C1100 (helix 37) and A1167 (helix 40), of 16S rRNA. The side chain of R69 interacts with both the phosphate group of C1100 and the carboxyl-terminus of Y71. The side chain of Y71 engages in a stacking interaction with base A1167. The interacting residues and nucleotides are shown as stick models. Hydrogen bonds are indicated by dashed lines. **b,** Structural comparisons of the inter-subunit bridge B2a and the decoding centre between the hibernating ribosome•RNase I complex (this study) and a classical state of the 70S ribosome (PDB 8SYL). Nucleotides showing notable conformational changes involved in bridge B2a formation or decoding centres are shown as stick models, along with the interacting amino acid residue (E68) from HPF. The N, O, and P-atoms are shown in blue, red, and orange, respectively.

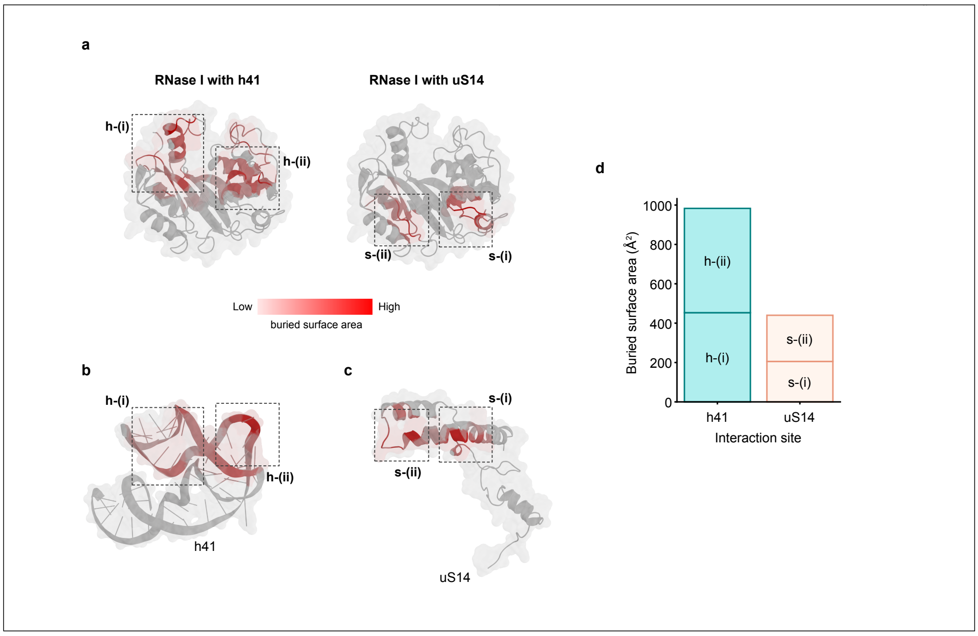

**Extended Data Fig. 6 | Buried surface area (BSA) of interaction sites of the hibernating ribosome•RNase I complex. (related to Fig. 3–d)**

**a,** Colour coding of the RNase I surface structure based on BSA values predicted using PISA for interaction with h41 (left) and uS14 (right). Residues with higher BSA values are indicated in darker red. RNase I is viewed from the bottom of the arch, as shown in Fig. 3c (right). **b-c,** Colour coding of h41 (b) or uS14 (c) surface structures based on BSA values predicted by PISA for interactions with RNase I. Colour maps are shared with (a). The views of h41 and uS14 are the same as those shown in Fig. 2c (right). **d,** The sums of BSA for each interaction region. The name of each interaction region shown in b-d corresponds to that shown in Fig. 3.

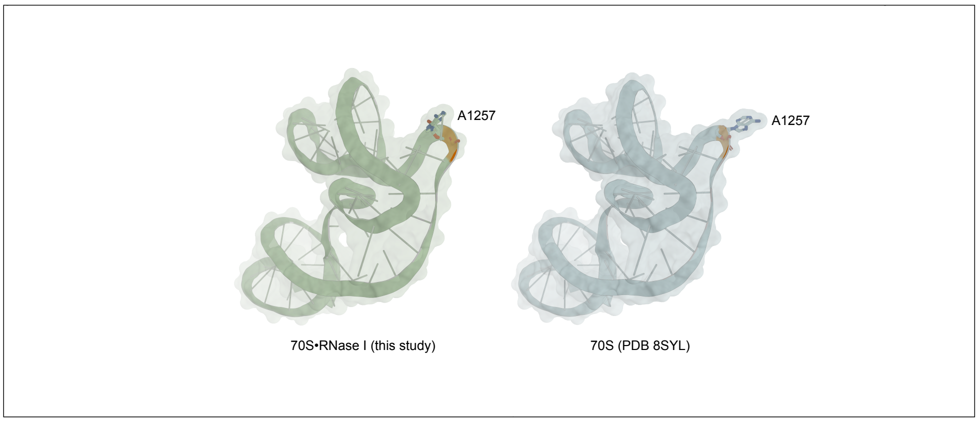

**Extended Data Fig. 7 | Conformational change of A1257**

Structural comparison of the base of A1257 in the h41 stem-loop between the hibernating ribosome•RNase I complex (this study) and a typical 70S ribosome (PDB 8SYL). h41 in our structure as well as in the 8SYL structure is represented by ribbon models with transparent surfaces and light green (the same as Fig. 2b) and whitish blue (the same as Fig. 2e) colours, respectively. Both A1257 nucleotides are represented by stick models, and the atoms of N, O, and P are shown in blue, red, and orange, respectively.

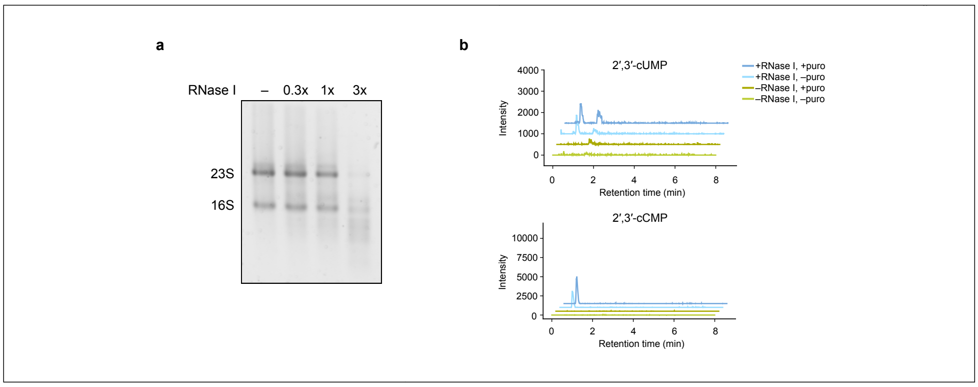

**Extended Data Fig. 8 | RNase I degrades rRNA within each subunit more efficiently than within 70S ribosomes.**

**a,** RNase I was added at the indicated molar ratio to 10 pmol of 70S ribosomes and incubated at 37 °C for 30 min. rRNA was prepared and analyzed by electrophoresis on a denaturing agarose gel. **b,** RNase I-mediated RNA cleavage in cells upon ribosome dissociation (related to Fig. 4b). Δ*rna* cells carrying either an empty vector or a plasmid expressing RNase I were treated with or without puromycin (puro), and the peaks of 2′,3′-cUMP and 2′,3′-cCMP in the cell extracts were detected by LC-MS. Extracted ion chromatograms for 2′,3′-cUMP (upper; *m/z* 307.0326 $[M+H]^+$) and for 2′,3′-cCMP (lower; *m/z* 306.0486 $[M+H]^+$) are shown.

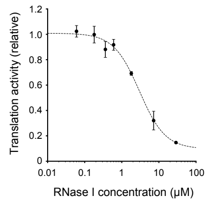

**Extended Data Fig. 9 | RNase I inhibits translation in a concentration-dependent manner.**

GFP was synthesized using the PURE system in the presence of various concentrations of RNase I. Translation activity was measured as the relative GFP fluorescence intensity. A sigmoidal curve was fitted to the data using the Hill equation, and the resulting fit is shown as a dashed line. The Hill coefficient was calculated to be 1.2, suggesting a typical 1:1 binding mode between the inhibitor and the ribosome. The $IC_{50}$ value was calculated to be 3.0 μM.

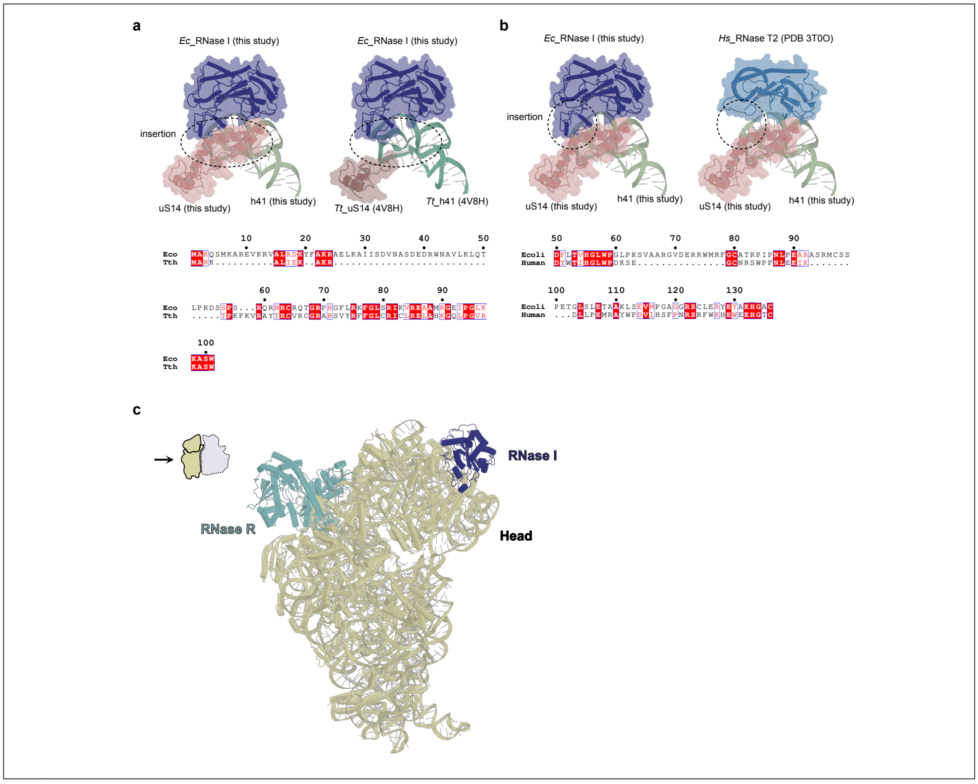

**Extended Data Fig. 10 | Structural features of the ribosomal protein uS14 and RNase I contributing to specific ribosome interaction, and comparison with other ribonucleases.**

**a,** (upper) uS14 and h41 of the structure of *T. thermophilus* 70S ribosome (PDB 4V8H[1]) are superimposed onto those of our structure shown in Fig. 3b. Our structure and the superimposed structure are shown on the left and right sides, respectively. (lower) Pairwise sequence alignment of uS14 from *E. coli* and *T. themophilus*. **b,** (upper) The structure of human RNase T2 (PDB 3T0O[2]) are superimposed onto our structure of RNase I shown in Fig. 3b. Our structure and superposed structure are shown on the left and right sides, respectively. (lower) Pairwise sequence alignment of *E. coli* RNase I and human RNase T2. **c,** RNase R from the structure of 30S•RNase R (PDB 8CDU[3]) superimposed on our structure shown in Fig. 3a. The 30S subunit, RNase I, and RNase R are shown in light yellow, dark blue, and light blue, respectively. Each chain is represented by a ribbon model (**a**–**c**). The transparent surfaces of *E. coli* RNase I, *E. coli* uS14, *T. thermophilus* uS14, *H. sapiens* RNase T2 are overlaid on the respective ribbon models (**a**, **b**). *E. coli* RNase I, human RNase T2, *T. thermophilus* uS14, and h41 are abbreviated as *Ec*_RNase I, *Hs*_RNase T2, *Tt*_ uS14, and *Tt*_h41, respectively.

### Extended Data Table 1 | Cryo-EM data collection, refinement and validation statistics

| | RNase I•70S hibernating complex<br>(EMD-60747)<br>(PDB 9IOT) |
|---|---|
| **Data collection and processing** | |
| Magnification | 80460x (calibrated) |
| Voltage (kV) | 300 |
| Electron exposure (e–/Å$^2$) | 50 |
| Defocus range (μm) | -0.65 to -2 |
| Pixel size (Å) | 1.21 |
| Symmetry imposed | C1 |
| Initial particle images (no.) | 1,501,256 |
| Final particle images (no.) | 50,955 |
| Map resolution (Å) | 2.8 |
| FSC threshold | 0.143 |
| **Refinement** | |
| Initial model used (PDB code) | 8SYL |
| Model resolution (Å) | 3.1 |
| FSC threshold | 0.5 |
| Map sharpening *B* factor (Å$^2$) | -39.8 |
| Model composition | |
| Non-hydrogen atoms | 239,377 |
| Protein residues | 6,062 |
| RNA nucleotides | 4,495 |
| Ligands | 477 |
| *B* factors (Å$^2$) | |
| Protein residues | 54.91 |
| RNA nucleotides | 54.72 |
| Ligands | 50.89 |
| Water molecules | 18.53 |
| R.m.s. deviations | |
| Bond lengths (Å) | 0.013 |
| Bond angles (°) | 1.973 |
| Validation | |
| MolProbity score | 2.48 |
| Clashscore | 3.47 |
| Rotamer outliers | 9.90 |
| Cβ outliers | 0.93 |
| Ramachandran plot | |
| Favored (%) | 88.68 |
| Allowed (%) | 9.51 |
| Disallowed (%) | 1.82 |

**Extended Data Table 2 | Hydrogen bonds at each interface in the hibernating ribosome•RNase I complex.**

| Ribosome | Distance (Å) | RNase I |
|---|---|---|
| **16S rRNA** | | |
| A1257[ N6 ] | 3.15 | ALA 81[ O ] |
| G1278[ N1 ] | 3.48 | GLU 90[ OE1] |
| G1278[ N1 ] | 3.38 | GLU 90[ OE2] |
| G1278[ N2 ] | 3.21 | GLU 90[ OE1] |
| G1278[ N2 ] | 3.19 | GLU 90[ OE2] |
| C1262[ N3 ] | 2.98 | ARG 31[ NH1] |
| C1273[ O2 ] | 3.01 | ARG 31[ NH2] |
| C1273[ O2'] | 2.98 | ARG 31[ NH2] |
| C1262[ O2'] | 3.7 | ASN 32[ ND2] |
| A1274[ O3'] | 3.01 | ARG 33[ NH1] |
| A1275[ OP1] | 2.78 | ARG 33[ NH1] |
| A1274[ O3'] | 3.17 | ARG 33[ NH2] |
| A1274[ O2'] | 2.62 | ARG 33[ NH2] |
| A1257[ O2'] | 3.67 | ARG 73[ NH1] |
| G1276[ O6 ] | 3.81 | ARG 77[ NH1] |
| A1256[ O2'] | 2.62 | ARG 83[ NH1] |
| A1256[ O3'] | 2.86 | ARG 83[ NH1] |
| A1257[ OP2] | 2.93 | ARG 83[ NH1] |
| G1278[ O6 ] | 2.89 | ARG 83[ NH2] |
| G1278[ OP2] | 3.71 | ARG 92[ NH1] |
| G1278[ OP2] | 3.66 | ARG 92[ NH2] |
| C1277[ OP2] | 2.92 | ARG 92[ NH2] |
| C1277[ OP2] | 3.05 | SER 94[ OG ] |
| G1272[ O2'] | 3.85 | ASN 233[ ND2] |
| **uS14** | | |
| ASN 35[ ND2] | 3.72 | GLY 232[ O ] |
| ASP 18[ O ] | 3.33 | GLY 69[ N ] |
| ASP 33[ OD2] | 2.8 | HIS 230[ ND1] |

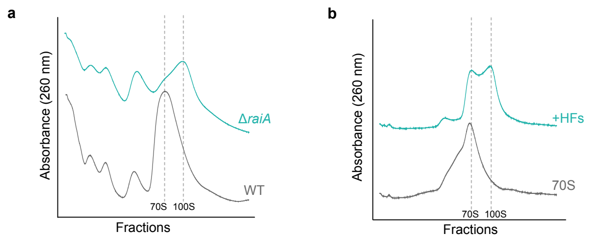

**Supplementary Fig. 1 | Confirmation of 100S ribosome formation by the sucrose density gradient centrifugation. a,** Profiles of ribosomes prepared from the wild-type strain in the exponential phase and from the RaiA-deficient (Δ*raiA*) strain cultivated in the amino acid starvation for 24 h. **b,** Profiles of 70S ribosomes and that with HFs (100S ribosomes). The sucrose density gradient centrifugation (SDGC) analysis was prepared as follows: (a) The cell pellets were resuspended in 1 mL of the ribosome gradient buffer (20 mM Tris-HCl (pH 7.5), 60 mM $NH_4Cl$, 7.5 mM $Mg(OAc)_2$, 0.5 mM EDTA, 6 mM β-mercaptoethanol) and disrupted by shaking for 5 min with 300 mg of zirconium silicate beads in a homogenizer (ShakeMaster Neo, Bio Medical Science Inc., Japan). The sample was then centrifuged for 10 min at 18,500 x g and 4 °C, and the resultant supernatant was collected. The absorbance at 260 nm ($A_{260}$) for lysates was determined (by Nanodrop), and 300 µL of normalized lysate was loaded on a 10% to 40% sucrose gradient in the ribosome gradient buffer. (b) After 3 h of centrifugation at 35,000 rpm and 4 °C using a Beckman SW41 rotor, the sample was collected with a BioComp gradient station and absorbance at 260 nm was continuously monitored. (b) The *in vitro* reconstituted hibernating ribosomes were prepared by incubating 23 pmol of 70S ribosomes with 240 pmol of RMF and 830 pmol of HPF in the reaction buffer (20 mM Tris-HCl (pH 8.0), 100 mM $NH_4Cl$, 10 mM $MgCl_2$) for 30 min at 37 °C. The profile of the resulting mixture was analyzed by the SDGC as above.

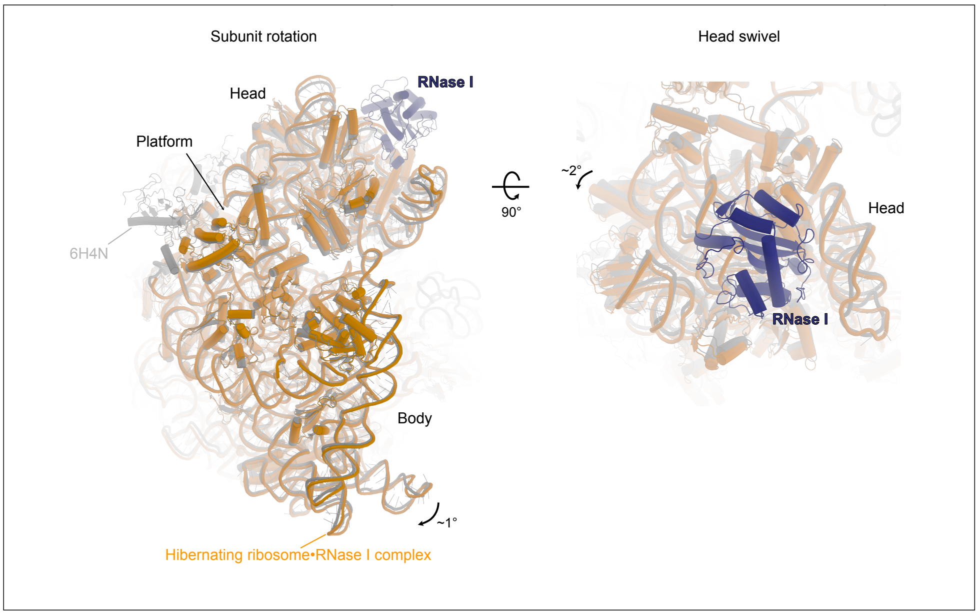

**Supplementary Fig. 2 | Conformational changes of the hibernating ribosome•RNase I complex.** In the hibernating ribosome•RNase I complex (ribosome in orange, RNase I in blue), the 30S subunit rotates clockwise by ~1° and the head domain swivels by ~2° compared to the hibernating ribosome lacking RNase I (gray; 6H4N).

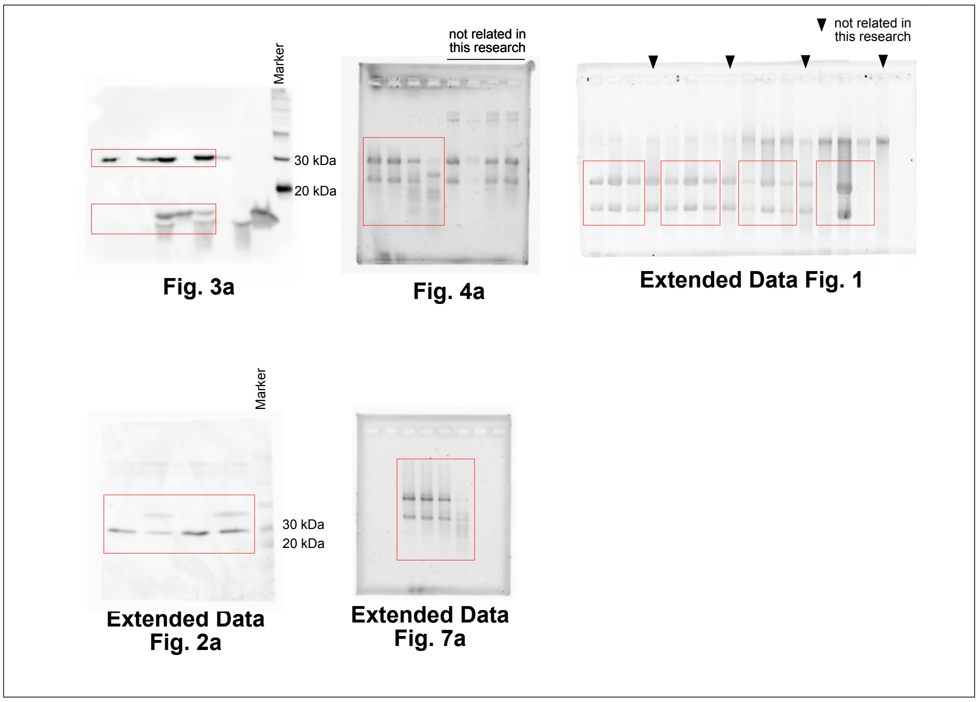

**Supplementary Fig. 3 | Source Data.** Uncropped western blots and gels used in Fig. 3a and 4a, and Extended Data Fig. 1, 2a, and 7a.

### Supplementary Table 1 | Strains and plasmids used in this study.

| Name | Description | Source |
|---|---|---|
| **Strains** | | |
| BW25113 | *lacI*$^{q}$ *rrnB*$_{T14}$Δ*lacZ*$_{WJ16}$ *hsdR514*Δ*araBAD*$_{AH33}$ Δ*rhaBAD*$_{LD78}$ | NBRP, Japan |
| Δ*rna* | *lacI*$^{q}$ *rrnB*$_{T14}$Δ*lacZ*$_{WJ16}$ *hsdR514*Δ*araBAD*$_{AH33}$ Δ*rhaBAD*$_{LD78}$ *rna*::Km$^{R}$ | NBRP, Japan |
| Δ*raiA* | *lacI*$^{q}$ *rrnB*$_{T14}$Δ*lacZ*$_{WJ16}$ *hsdR514*Δ*araBAD*$_{AH33}$ Δ*rhaBAD*$_{LD78}$ *raiA*::Km$^{R}$ | NBRP, Japan |
| Δ*rmf* | *lacI*$^{q}$ *rrnB*$_{T14}$Δ*lacZ*$_{WJ16}$ *hsdR514*Δ*araBAD*$_{AH33}$ Δ*rhaBAD*$_{LD78}$ *rmf*::Km$^{R}$ | NBRP, Japan |
| Δ*hpf* | *lacI*$^{q}$ *rrnB*$_{T14}$Δ*lacZ*$_{WJ16}$ *hsdR514*Δ*araBAD*$_{AH33}$ Δ*rhaBAD*$_{LD78}$ *hpf*::Km$^{R}$ | NBRP, Japan |
| Δ*raiA*Δ*rna* | *lacI*$^{q}$ *rrnB*$_{T14}$Δ*lacZ*$_{WJ16}$ *hsdR514*Δ*araBAD*$_{AH33}$ Δ*rhaBAD*$_{LD78}$ *raiA rna*::Km$^{R}$ | This study |
| Δ*raiA*Δ*hpf*Δ*rmf*Δ*rna* | *lacI*$^{q}$ *rrnB*$_{T14}$Δ*lacZ*$_{WJ16}$ *hsdR514*Δ*araBAD*$_{AH33}$ Δ*rhaBAD*$_{LD78}$ *raiA hpf rmf rna*::Km$^{R}$ | This study |
| W3110 | K-12 (F$^{-}$ λ$^{-}$) | NCBI |
| BLR (DE3) | F$^{-}$ *ompT hsdS*$_{B}$(r$_{B}^{-}$ m$_{B}^{-}$) *gal lac ile dcm* Δ(*srl-recA*)::Tn10 (tet$^{R}$)(DE3) | Novagen |
| **Plasmids** | | |
| pUC18 | A plasmid with an ampicillin resistance marker | TaKaRa Bio |
| pUC18-*rna* | Expressing RNase I in *E. coli* | This study |
| pET24d | A plasmid with a kanamycin resistance marker for inducible expression of protein under T7 promoter | Merck |
| pET-*rna* | Expressing His-tagged RNase I under T7 promoter | This study |
| pET-*rna*_mt | Expressing His-tagged RNase I (H55F/H133F) under T7 promoter | This study |
| pET-*rmf* | Expressing His-tagged RMF under T7 promoter | This study |
| pET-*hpf* | Expressing His-tagged HPF under T7 promoter | This study |
| pET-*gfp* | Expressing GFP under T7 promoter | This study |
| 707-FLPe | Encoding flippase and tetracycline resistance genes | Gene Bridges |
